# Molecular mechanism of thyroxine transport by monocarboxylate transporters

**DOI:** 10.1101/2024.10.17.618737

**Authors:** Matteo Tassinari, Giorgia Tanzi, Francesco Maggiore, Stefan Groeneweg, Ferdy S. van Geest, Matthijs Freund, Christiaan J. Stavast, Irene Boniardi, Sebastiano Pasqualato, W. Edward Visser, Francesca Coscia

## Abstract

Thyroid hormones (the common name for prohormone thyroxine and the bioactive form triiodothyronine) control major developmental and metabolic processes. Release of thyroid hormones from the thyroid gland into the bloodstream and their transport into target cells is facilitated by plasma membrane transporters, of which monocarboxylate transporter (MCT)8 and the highly homologous MCT10 are most important. Patients with MCT8 mutations suffer from a severe neurodevelopmental and metabolic disorder, however, the molecular mechanism underlying thyroid hormone transport is unknown. Using cryogenic-sample electron microscopy (cryo-EM), we determined the ligand-free and thyroxine-bound human MCT8 structures in the outward-open state and the thyroxine-bound human MCT10 in the inward-facing state. Our structural analysis revealed a network of conserved gate residues involved in conformational changes upon thyroxine binding, triggering ligand release on the opposite compartment. We then determined the structure of a folded, but inactive patient-derived MCT8 mutant, indicating a subtle conformational change which explains its reduced transport activity. In addition, we determined the structure of MCT8 bound to its inhibitor silychristin, revealing an interaction with residues essential to drive transition to the inward-facing state, thereby locking the protein in the outward-facing state. This study provides molecular and structural insights into normal and disordered thyroid hormone transport.

## Introduction

Thyroid hormones are critical for normal development and metabolism in virtually all cells^1^. Plasma membrane transporters facilitate transport of thyroid hormones across the plasma membrane, thereby governing intracellular thyroid hormone concentrations^2,3^. Monocarboxylate transporter (MCT) 8 and MCT10 facilitate transcellular transport of the prohormone T4 (3,5,3′,5′-tetraiodothyronine or thyroxine) and of the bioactive T3 (3,5,3′- triiodothyronine)^4,5^. MCT8 is the most proficient thyroid hormone transporter and facilitates release of thyroid hormones from the thyroid and transport across the blood-brain barrier and into neuronal cells^6^. In vitro, MCT8 is specifically inhibited by the flavonolignan silychristin^5,7^. Loss-of-function (LoF) mutations in the X-chromosomal *SLC16A2* gene, encoding MCT8, cause severe intellectual and motor disability due to cerebral hypothyroidism and clinical features such as underweight and tachycardia secondary to chronic peripheral thyrotoxicosis, with ∼30% dying in childhood ^8^. Survival rate and the severity of disease features is strongly determined by the functional impact of the LoF mutation *(Groeneweg/van Geest, Nat Comm, in principle accepted)*. Treatment with the T3 analog TRIAC, which is able to enter cells independent of MCT8, is able to alleviate peripheral thyrotoxicosis symptoms^9,10^. Among monocarboxylate transporters, MCT10 is the most similar to MCT8, with 49% sequence identity and the only other thyroid hormone transporter in this protein class^11^. However, the physiological function of MCT10 is currently unclear. Despite the critical role of these plasma membrane transporters, their substrate binding and transport mechanisms have been elusive^12,13^.

Here, utilizing cryogenic-sample Electron Microscopy (cryo-EM) we determined five structures describing different states of the transport cycle: human MCT8 in the ligand-free, thyroxine-bound, silychristin-bound, a patient-derived D424N (D498 in the long isoform) in the inward-open conformation and thyroxine-bound human MCT10 in the outward facing conformation. Our structural comparison combined with biophysics binding and cellular transport assays provides insight in the normal and disordered mechanisms of action of these two homologous thyroid hormone transporters^11^.

## Results

### Thyroid hormone binding and transport

To study thyroid hormone binding and transport, we genetically fused an N-terminal FLAG tag to both targets. First, we verified that upon overexpression of FLAG-MCT8 (Figure 1a,b) in HeLa cells, T4 added in the cellular medium is efficiently internalized, while addition of the inhibitor silychristin reduces MCT8-mediated transport (Figure 1a,b).

**Figure 1:**
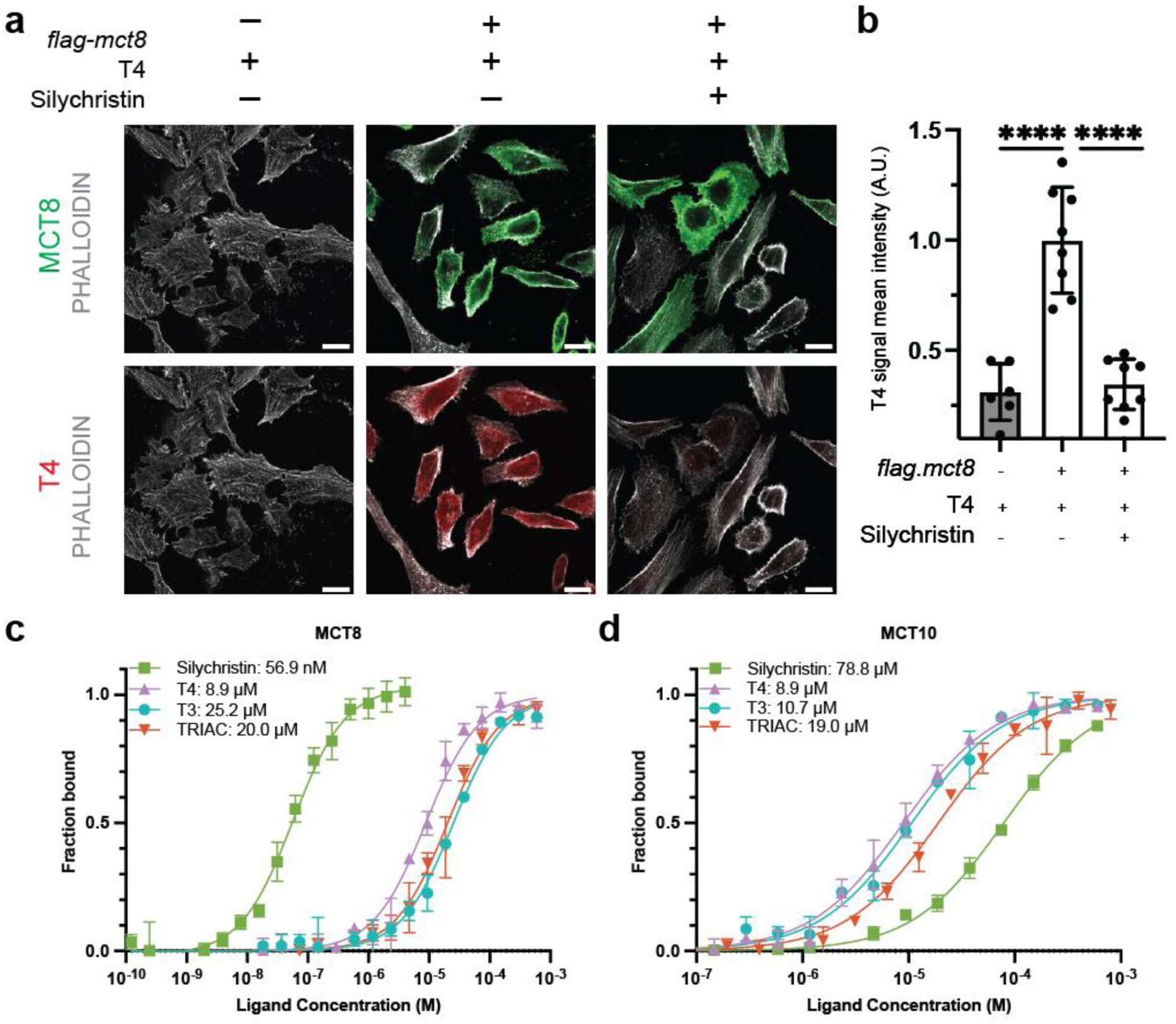
T4 transport and inhibition by silychristin by MCT8. **a.** Representative confocal images of three independent experiments of Hela cells overexpressing FLAG-MCT8, exposed to 10 µM T4 in absence or in presence of 10 µM silychristin. Detection of T4 and MCT8 is performed by immunofluorescence. Scale bar: 20µm. P-values based on ordinary one-way ANOVA (**** p< 0.0001). **b.** Quantification of T4 internalization by HeLa cells overexpressing FLAG-MCT8. Normalisation was performed on overexpressed MCT8+T4 Hela cells **c-d**. Ligand binding curves and relative affinities purified for MCT8 and MCT10 measured *in vitro* by label-free microscale thermophoresis (MST).

We then isolated overexpressed FLAG-MCT8 and FLAG-MCT10 in detergent, which are purified as a monodisperse peak in size-exclusion chromatography (Supplementary Figure 1 a,b). Although higher order species are visible by SDS-PAGE analysis (Supplementary Figure c,d) both proteins are monomeric and folded within the detergent micelle, as further verified by cryo-EM analysis (Supplementary Figure 1 e,f).

We measured affinity of purified MCT8 and MCT10 for thyroid hormones *in vitro*, using label-free microscale thermophoresis (Figure 1c,d and Supplementary Figure 1). FLAG- MCT8 showed a Kd of 8.9 µM for T4, of 25.2 µM for T3 and 56.9 nM for silychristin, the latter being confirmed by isothermal titration calorimetry (ITC) measuring a Kd of 44.5 nM (Supplementary Figure 2). FLAG-MCT10 showed a Kd of 8.9 µM for T4, of 10.7 µM for T3 while binding to silychristin is beyond 75 µM. These findings are in line with previous data^4,5,6^ and confirmed preserved activity of purified recombinant FLAG-tagged MCT constructs.

### Structures of MCT8 and MCT10 along the T4 transport cycle

In the present study, we numbered the residues according to the short MCT8 isoform; to convert the number to the long MCT8 isoform (which has been used in the past to describe patient mutations and in homology models), 74 amino acids should be added. Our first attempts for structural characterization of MCT8 and MCT10 by cryo-EM, both in detergents and amphipols, were not successful as we experienced low-resolution reconstructions or severe preferred orientation within the EM grid, respectively, precluding accurate substrate and protein model building. We then engineered both MCT8 and MCT10 with a C-terminal ALFA tag (see constructs paragraph in the methods section) and reconstituted a complex with an anti-ALFA nanobody (NbALFA in Figure 2a) and in presence of amphipols (Supplementary Figure 3). This strategy considerably improved the orientation distribution (Supplementary Figure 4), enabling us to obtain cryo-EM maps of the ligand-free MCT8 in the outward-facing state (OFS), T4-bound MCT8 in the OFS and of T4-bound MCT10 in the inward-facing state (IFS) at the resolutions of 3.5 Å, 3.4 Å and 3.8 Å, respectively (Figure 2b, c, e). Both MCT8 and MCT10 display the classical bilobed architecture of the major facilitator superfamily (MFS), including a N-terminal domain (NTD) formed by transmembrane helices (TMH) 1 to 6 and C-terminal domain (CTD) formed by TMH 7-12^14,15,16^. Clear densities for the ligand T4 are visible in our cryo-EM maps of both MCT8 and MCT10 (Figure 2d and f). In all structures of MCT8, the CTD residues facing the solvent-exposed central protein gate Y339 (TMH7) and R371 (TMH8) are linked via an H-bond and a salt bridge through D424 (TMH10) (Supplementary Figure 5). The MCT8 pocket accommodates the two iodinated aromatic rings of T4 in an inner highly hydrophobic pocket, while the amino acidic moiety is positioned towards the outward wide opening (Figure 2h). The T4 carboxylic group is bound via a salt bridge to R371, while the amino group of the substrate points towards N119 on the NTD lobe, although not through direct interactions. The ligand-free and T4- bound MCT8 structures appear almost identical (RMSD 1.018 Å), but we observe a subtle conformational change where the TMH7 kinks. In particular, the F336-Y335 pair moves closer to the substrate, causing Y339 (along the same TMH7) to approach N119 (Cα-Cα distance from 13.8 to 11.2 Å) on the opposite NTD lobe, thereby partially occluding the gate (Figure 2gh and jk). In the T4-bound MCT10 structure (IFS) the CTD is stabilized by the same interactions as in MCT8 (Y311 on TMH7 is held to R343 on TMH8 via D396 on TMH10). T4 is rotated with respect to MCT8. Y307, Y311 and N89 (Y335, Y339 and N119 in MCT8) coordinate the carboxylic and amino group of T4 at about 3 Å distance. Hence, the Y311-N89 pair seals the gate (Cα-Cα distance 6.4 Å) on the outward facing compartment. Thus, T4 becomes accessible to the opposite compartment, where it is conveyed by concentration gradient (Figure 2i and l). Overall, the comparison among the structures of the highly conserved transporters in the OFS and IFS suggests that the transport of thyroxine in both MCT transporters is initiated upon sensing of the substrate mostly on the CTD lobe. The latter is tightly held by the central residue D424, allowing a concerted closure of CTD and NTD onto the amino acid group of T4.

**Figure 2:**
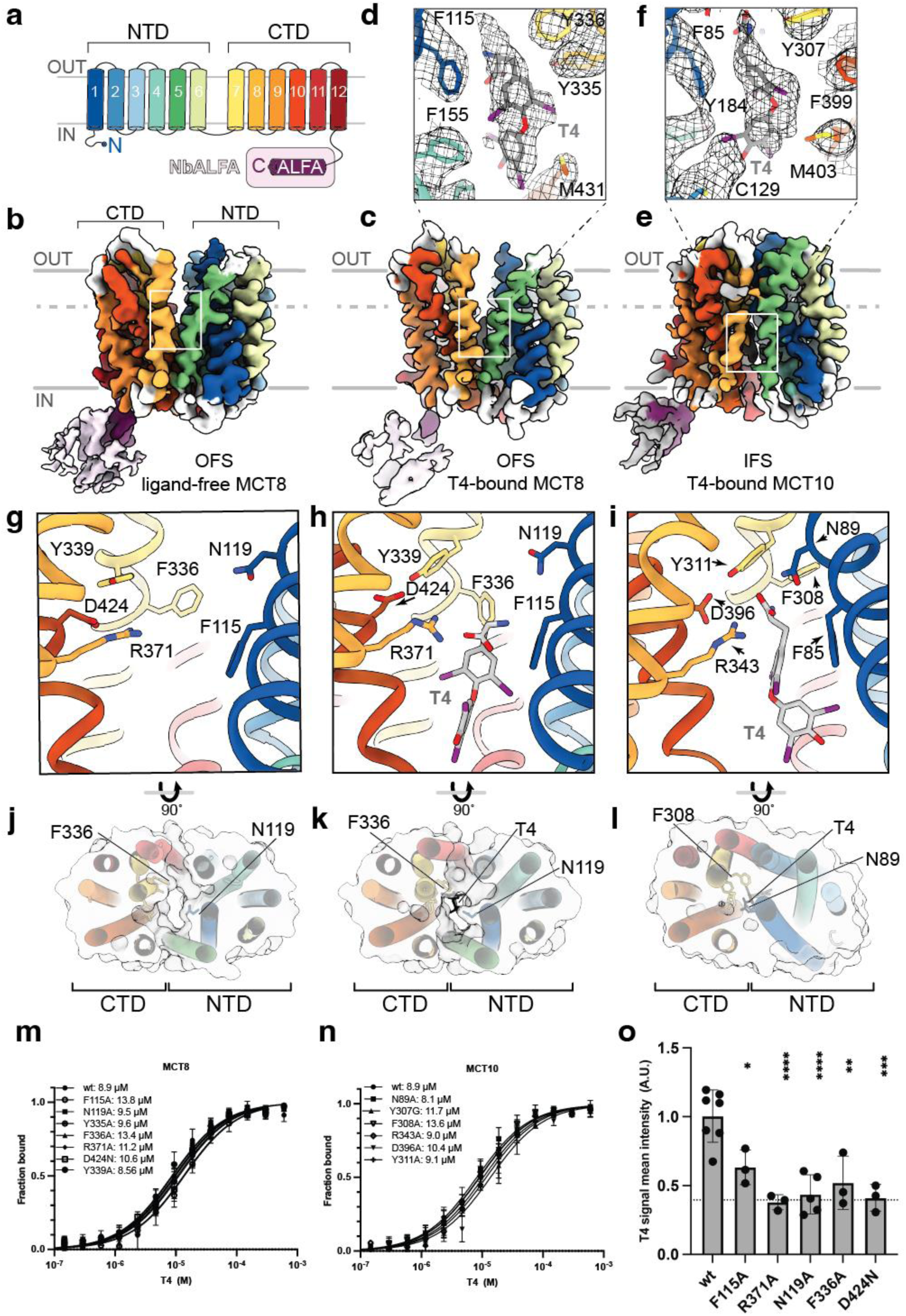
Structures of MCT8 and MCT10 in complex with T4. **a.** Schematic representation of monocarboxylate transmembrane helices 1-6 (N-terminal domain, NTD) and 7-12 (C-terminal domain, CTD). The construct was engineered with an ALFA tag at the C-terminus to form a complex with its cognate nanobody NbALFA. **b.** cryo-EM map of ligand-free MCT8 in the outward facing state (OFS) **c.** cryo-EM map of T4-bound MCT8 in the outward facing state (OFS) **d.** Closed-up view on the MCT8 ligand binding site with relative EM density **e.** cryo-EM map of T4-bound MCT10 in the inward facing state (IFS) **f.** Closed-up view on the ligand binding site with relative EM density **g-i.** Zoomed views on the ligand binding site corresponding to the white square in b,d,e panels, highlighting residues coordinating the ligand **j.-l.** Cross section of the surface of transporters (dotted line in panels b,d,e) indicating progressive closure of the gate, moving solvent accessibility of the substrate from the OUT to the IN compartment. To translate the position of residues to the long isoform of MCT8, 74 amino acids should be added. **m.** T4 binding curve of MCT8 WT and mutants measured by MST. **n.** T4 binding curve of MCT10 WT and mutants measured by MST. **o.** T4 transport efficiency measured by cell-based transport assay (immuno-detection) for overexpressed MCT8 mutants. Data are normalised on MCT8 WT +T4. Dotted line indicates negligible endogenous MCT8 levels. P-values based on ordinary one-way ANOVA (* p< 0.05, ** p< 0.01, *** p< 0.001, **** p< 0.0001). Quantification of T4 internalization by HeLa cells overexpressing FLAG-MCT8. Normalisation was performed on overexpressed MCT8+T4 Hela cells

To corroborate these observations, we measured the affinity for T4 and transport efficiency of MCT8 (and corresponding MCT10) variants in which we mutated residues participating in the binding to the hydrophobic moiety of T4 (F115) and to the amino acid group (R371, N119), F336 involved in the transition from OFS to IFS (Figure 2m,n,o) and D424 stabilising the CTD. Intriguingly, none of the single residues is essential for substrate binding, but mutation of N119/N89, D424/D396 and R371/R343 severely reduced T4 transport, i.e., and appear essential for the transition from OFS to IFS (Figure 2o). In conclusion, our comparison between these three structures allowed us to identify a set of key residues involved in mechanism of thyroxine transport by MCT8 and MCT10. Our findings are also in agreement with previous functional studies^17^.

### Structure of the MCT8 pathogenic variant D424N

Structural analysis of MCT8 and MCT10 in presence of T4 highlighted that D424 is a key residue not directly involved in substrate binding, but important for transport. Indeed, the patient-derived mutation c.1492G>A corresponds to the D424N variant, associated with clinical features at the severe end of the phenotypic spectrum of MCT8 deficiency^18^. To understand why this isosteric mutation affects T4 transport and to support the hypothesized mechanism of transport, we determined the architecture of MCT8-D424N mutant by cryo-EM, obtaining a map at an overall resolution of 4 Å (Figure 3a). Our model shows that the protein variant is in the OFS, with an overall folding similar to ligand-free MCT8 (RMSD 1.8 Å). Nevertheless, in MCT8-D424N TMH 7 and TMH10 are shifted away from the main gate axis by consequence of the weaker bond between Y339 and N424 (Figure 3b,c). Overall, the map suggests an increased flexibility of TMH and on average a wider opening of the outward-facing lobes (Figure 3b). As measured by MST and by visual inspection of the ligand binding pocket, the protein is still competent for T4 binding (Figure 2m). Nevertheless, the absence of the crucial D424, which holds together the CTD, hampers an efficient TMH7-initiated conformational change leading to the hormone transport. In conclusion, the structure of the pathogenic MCT8 D424N mutant confirms the mechanism of thyroid hormone transport derived by the WT structures and suggests a rationale for its defective activity.

**Figure 3:**
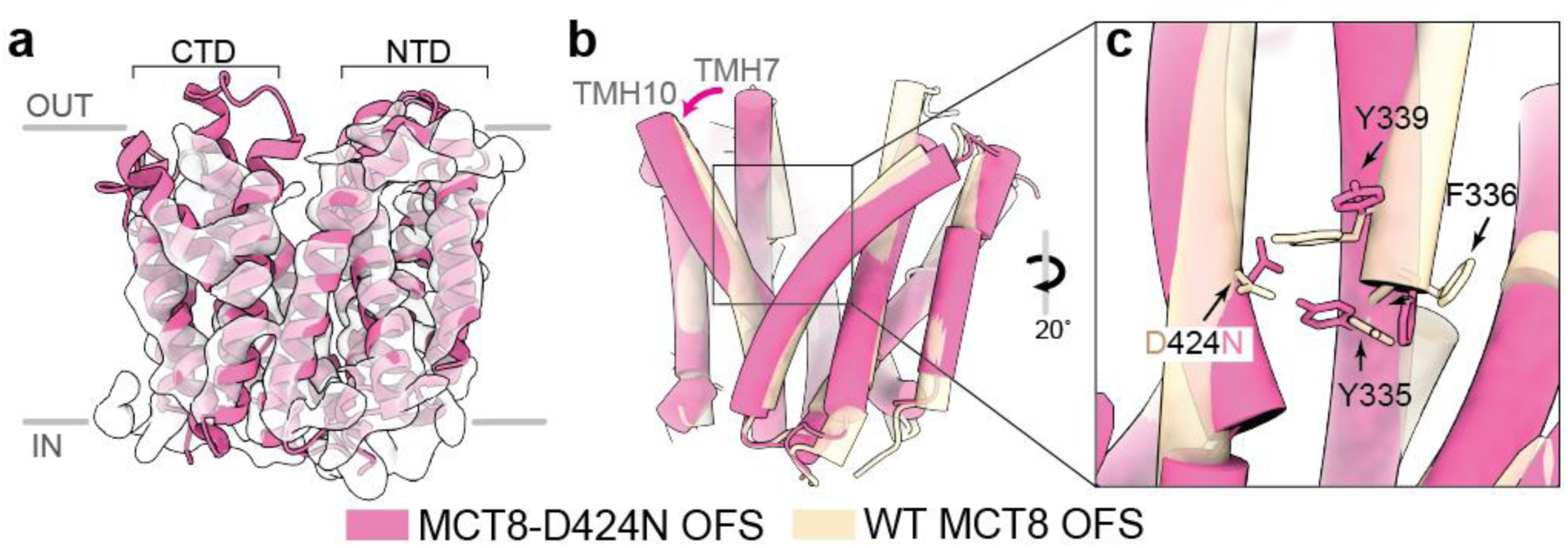
Structure of the patient-derived variant MCT8 D424N. **a.** cryo-EM map of MCT8 D424N (D498N) corresponding to an outward-facing state (OFS). **b-c.** Overlap between ligand-free WT MCT8 and MCT8 D424N (D498N), showing larger flexibility and lateral shift in TMH7 which is more loosely held to TMH10 by a weaker Y339-N424 (Y413-N498) interaction. To derive the position of residues in the long isoform of MCT8, 74 amino acids should be added (between brackets in the legend).

### Mechanism of MCT8 inhibition by silychristin

Having derived a mechanism of transport of T4 in MCT8 and MCT10, we set to investigate the mechanism of inhibition of MCT8 by determining its cryo-EM structure in complex with silychristin (Figure 4). Our 3.0 Å resolution cryoEM map allowed us to identify the binding pocket for the ligand within an OFS (Figure 4a,b). Silychristin is located in the same binding pocket of T4, but it is shifted towards the outward compartment with respect to the thyroid hormone. Superposition of the MCT8 structures bound to the inhibitor and to T4 revealed that the three aromatic moieties of silychristin do not overlap with the iodophenol rings of T4 (Figure 4c). Silychristin establishes close interactions with F336, R371 within the CTD and F115, N119 on the opposite NTD, by locking the protein in an apo-like conformation in which it is unable to undergo the conformational change necessary to initiate the transport (Figure 4f). To validate these structural findings, we performed MCT8 site-directed mutagenesis and measured the effects on silychristin binding and ability to inhibit T4 transport in cells (Figure 4d,e). Our data show that F115A abolishes binding and, hence, inhibition by silychristin in MCT8. MCT8 mutants R371A and F336A can still partially bind silychristin, but T4 transport is no longer inhibited by the binding of silychristin. A similar effect on silychristin binding is observed by preincubating MCT8 with 100 µM T4, indicating that when T4 occupies the gate first and partially occludes it, silychristin can no longer efficiently compete with its binding *in vitro*. To further investigate this aspect, we performed a competitive T4 binding experiment in presence of a large excess of the inhibitor. The binding curve indicates that if silychristin first occupies the gate, T4 affinity decreases from 8.9 to 20.6 µM (about 65%) but binding is not completely abolished. However, in presence of Silychristin MCT8 can no longer transport the ligand as F336 and N119 are stabilized by direct interaction with the inhibitor. In conclusion, our structural and functional data on the MCT8-silychristin complex indicate that the inhibitor does not completely prevent thyroid hormone binding but abolishes MCT8-mediated T4 transport by keeping the protein in an OFS.

**Figure 4:**
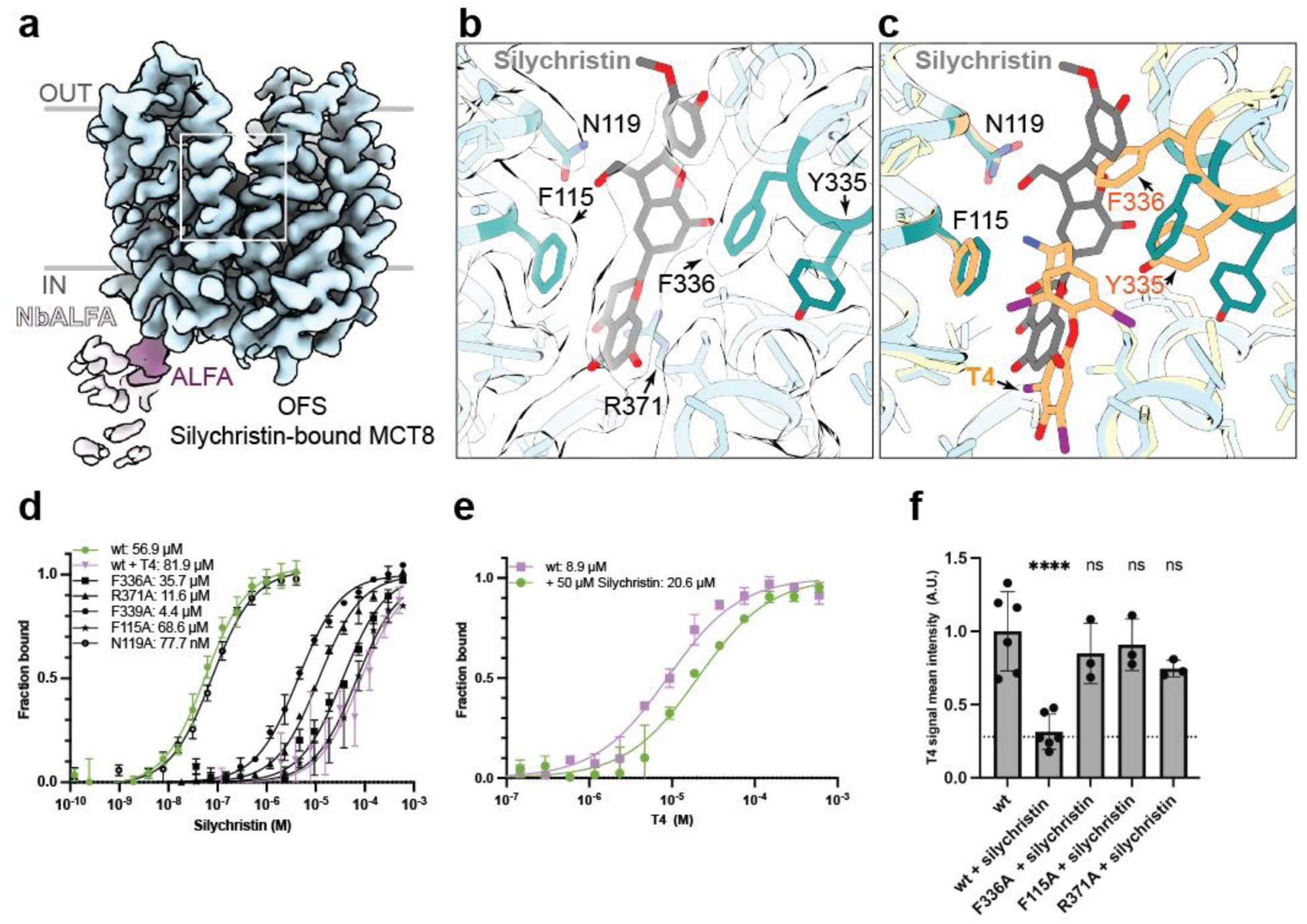
Structure of MCT8 bound to its inhibitor silychristin. **a.** cryo-EM map of silychristin-bound MCT8 in the outward facing state (OFS) **b.** Close-up view on the ligand binding site with relative EM density in white, showing interaction with T4 binding and transport residues, F336 (F410), N119 (N193), F115 (F189), R371 (R445). **c.** Overlap between T4-bound (gold) and silychristin-bound MCT8 in the OFS, showing that silychristin locks the protein in the OFS preventing movement of F336 and N119 (N193) necessary for transport. **d.** Silychristin binding curves of WT MCT8 (green), mutants (black) and WT preincubated with T4 (violet) obtained by MST. The curves show that the inhibitor binding is reduced when mutating F336 (F410) and R371 (R445) and Y339 (Y413), or when the gate is being already occupied by T4, but not when modifying N119 (N193). **e**. MCT8 T4 binding profile (MST) when the sample was preicubated with 50 µM silychristin, showing reduced ability to bind the natural substrate in presence of the inhibitor **f.** Silychristin inhibition of MCT8-mediated T4 transport measured by cell-based transport assay for WT and different MCT8 mutants. Values for each mutant were normalised with respect to its not-inhibited conditions (with T4 and without silychristin) (Figure 2o). Dotted line indicates negligible endogenous MCT8 levels. P-values based on ordinary one-way ANOVA (**** p< 0.0001). Quantification of T4 internalization by HeLa cells overexpressing FLAG-MCT8. Normalisation was performed on overexpressed MCT8+T4 Hela cells. To derive the position of residues in the long isoform of MCT8, 74 amino acids should be added (between brackets in the legend).

## Discussion

Thyroid hormone transporters are cellular gatekeepers for the regulation of intracellular thyroid hormone concentrations, orchestrating developmental and metabolic events in many tissues. In this work, we determined five cryo-EM structures of human MCT8 and MCT10 as exemplary thyroid hormone transporters and combined biophysical and functional studies to dissect binding and mechanism of transport of thyroid hormone as well as the molecular basis of its inhibition by silychristin.

A key finding suggested by our structures of MCT8 was that upon substrate binding, a subtle conformational change in TMH7 (F336) initiates a continuous movement of CTD TMHs, stabilized by the pivotal residue D424, which progressively closes the gate onto the amino acidic group of the ligand together with the N119 on the opposite NTD lobe, thereby exposing the hormone to the opposite compartment (Figure 5). T4 is bound to MCT8 via both hydrophobic interactions coordinating its aromatic moieties and with the aminoacidic group, essential to drive its transport via protein-ligand interactions (e.g., with Y335, N119, Y339). The bioactive hormone T3 (bearing a mono-iodinated phenol ring) binds with lower affinity to MCT8 (Figure 1c), but has comparable transport efficiency as T4 ^4,19^. We hypothesize that differential binding and transport of these very similar hormones is due to higher retention of T4 in the binding pocket, given its larger binding surface, which is then less efficiently released by the protein upon exposure to the gradient concentration. Previous reports have highlighted that di-iodo-tyrosine (DIT) is also a substrate for MCT8 ^20^, while the T3 analogue TRIAC is not transported by MCT8^21^. These findings are in agreement with the proposed mechanism of transport, as DIT lacks a di-iodo-ring with respect to T4, but bears the amino acidic group necessary for transport. Conversely, TRIAC bears the same T3 aromatic moiety, but lacks the amino group. Indeed, TRIAC binds MCT8 similarly as T3 (Figure 1c-d) but it is not transported by MCT8, potentially acting as a mild MCT8 inhibitor. This hypothesis is in agreement with cis-inhibitory studies showing that TRIAC inhibits MCT8-mediated T3 transport^19^.

**Figure 5:**
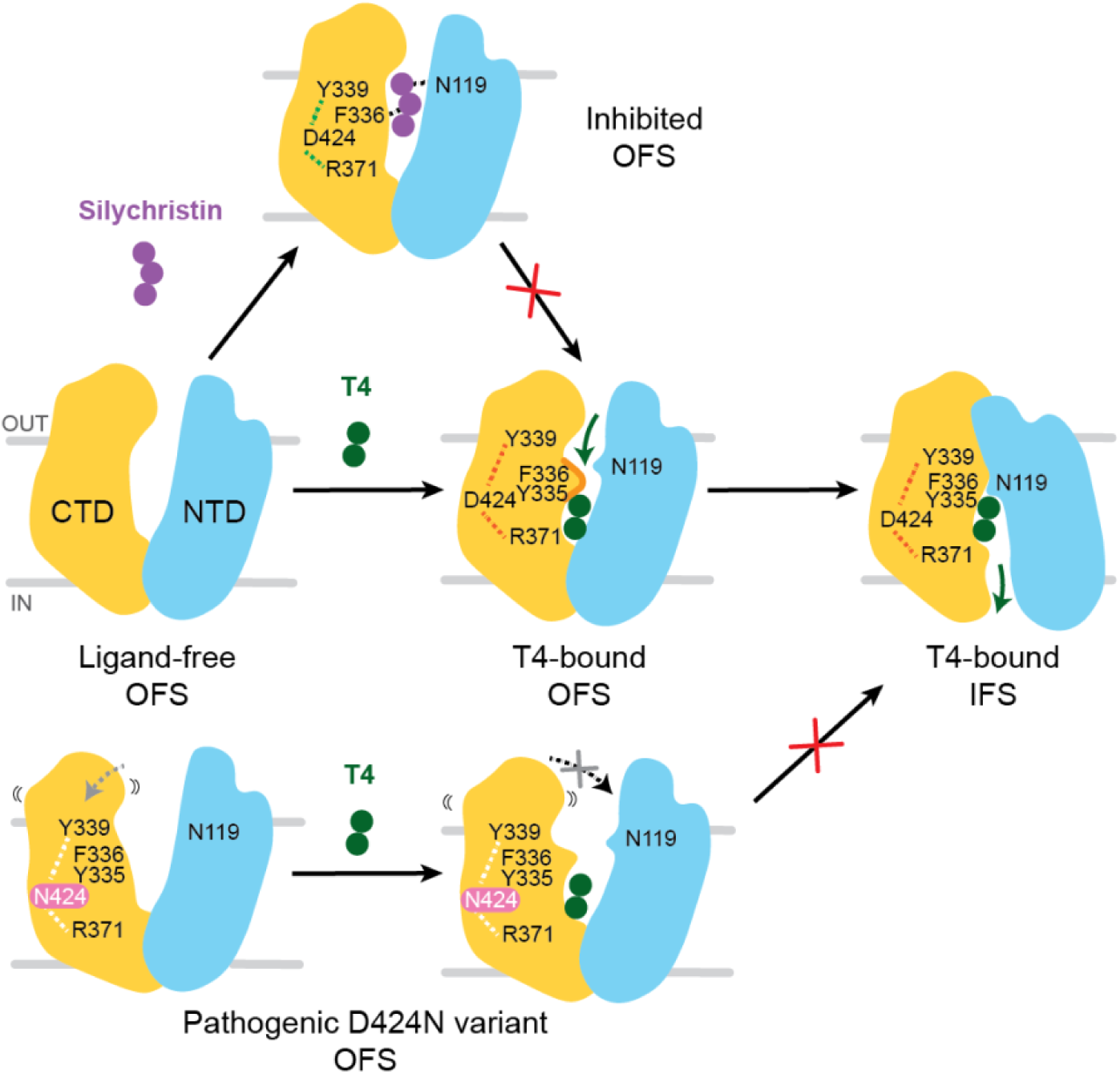
Proposed model of the mechanism of MCT8-mediated thyroid hormone transport and its inhibition. Schematic illustration of MCT8 conformations during the thyroid hormone transport cycle. The states represent experimentally obtained conformations in this study (with the T4-bound IFS cartoon derived from T4-bound MCT10).

By comparing the structures of MCT8-T4 and the proton-coupled MCT1 in the OFS bound to its substrate lactate^22^ (PDB ID 6LZ0), we observe that the carboxylic group is in a similar position with respect to the lactate coordinated by the conserved residue R371 (R313 in MCT1). However, D309, driving proton coupled transport is a serine (S367) in MCT8, while D424 is replaced by a phenylalanine. Moreover, the MCT1 pocket, tailored to lactate, is occupied by large hydrophobic residues thereby making it incompatible with thyroid hormone binding. R371 is highly conserved in the pocket across human MCTs (Supplementary Figure 6) and likely a defining residue coordinating the negatively charged substrates of this class of major facilitator superfamily (MFS), in accordance with previously reported homology models^13,12^. Our structures confirm previous functional data, indicating that the R371 side-chain interacts with substrate molecules and is a target for chemical modification^12,13,17^. Conversely, the Y339 and D424 pair is exclusively present in the thyroid hormone transporters MCT8 and MCT10 (Supplementary Figure 6b)^11^. Previous studies also suggested an important role for H118 in substrate binding^23,12^. Here, we show that the H118 side-chain is indeed exposed to the solvent and is thus available for chemical modification by His-reactive diethyl pyrocarbonate, but does not directly interact with T4 in the currently determined conformational states. In this work, we also revealed the mode of binding and inhibition of MCT8 by silychristin, which coordinates both the NTD and CTD forcing MCT8 in a OFS. Interestingly, our binding studies indicate that T4 is still weakly binding MCT8 in presence of silychristin, but its transport can no longer occur given the large number of interactions of the inhibitor, with a Kd considerably lower with respect to T4.

Overall, the mechanism of transport and involvement of a set of key residues was confirmed by the structure of the pathogenic mutant D424N, which is still able to bind T4 but its transport efficacy is highly reduced due to larger flexibility of the CTD which prevents an efficient adaptive closure of the gate (Figure 5).

In conclusion, combining structural findings with functional assays we dissected the principles of binding and transport of thyroid hormones to monocarboxylate transporters and identified essential residues regulating access to two opposite membrane-separated compartments. In addition, our ALFA-tag fusion allowed us to readily determine MCT8 structures of pathogenic variants. This opens the way to a faster determination of MCTs variants architecture, and together with the WT structures, to build a structure-educated interpretation of pathogenic variants. Taken together, our results contribute to further understanding of the principles and substrate specificity of monocarboxylate transporters and their mechanisms of transport.

## Methods

### Constructs

N-terminally FLAG-tagged pcDNA3-hMCT8 was previously generated^24^. A FLAG-tag was subcloned at the N-terminus of the previously generated pcDNA3.1-MCT10 plasmid using HindIII and Xbal restriction sites after polymerase chain reaction amplification of WT MCT10 cDNA using forward primers containing the FLAG-tag sequence (Supplementary figure 7)^5^. We numbered the residues according to the short MCT8 isoform; to convert the number to the long MCT8 isoform (NM_006517.3), which has been used in the past to describe patient mutations and in homology models, 74 amino acids should be added. The position of the mutations is indicated using the NM_006517.3 reference sequence and uses +1 as the A of the ATG translation initiation codon of the long MCT8 isoform, with the initiation codon as codon 1. DNA sequencing confirmed the presence of the introduced mutations and the absence of unintended mutations.

For cryo-EM structural studies of MCT8, both in its unbound form and in complex with L-thyroxine or silychristin, an ALFA tag (SRLEEELRRRLTEP) was introduced at position P505. In the MCT8 D424N variant, the ALFA tag was placed at F512. For MCT10 bound to L-thyroxine, the tag was fused at P470 (Supplementary figure 8). The following C-terminal residues were deleted.

For MCT8 and MCT10 site-directed mutagenesis, Q5 DNA polymerase (NEB) and Hi-Fi DNA assembly (NEB) were used.

### L-thyroxine internalization assay

HeLa cells were cultured in Dulbecco’s Modified Eagle’s Medium (DMEM, Euroclone, ECB7501LX10) supplemented with 10% (v/v) heat-inactivated fetal bovine serum (FBS) (Life Technologies, 10500064), 100 IU/ml penicillin–streptomycin (Euroclone, ECB3001D), and 2 mM L-glutamine (Euroclone, ECB3004D) at 37°C under atmospheric O2 and 5% CO2.

HeLa cells were plated in 24-well plates at a density of 50,000 cells per well and allowed to adhere for at least 24 hours. Cells were then transfected with MCT8 or MCT10 vectors (wild-type or mutants, see Table 1) using Lipofectamine 2000 (Invitrogen, 11668019).The medium containing the transfection reagents was replaced with fresh medium 5 hours post-transfection. The day after transfection, 10 µM L-thyroxine and/or 10 µM Silychristin were added to the medium for 20 minutes. The wells were washed three times with PBS, and cells were fixed with 4% PFA for 10 minutes at room temperature. Cells were then washed three times for at least 5 minutes in PBS before blocking for 1 hour at room temperature with PBS supplemented with 0.1% (v/v) Triton X-100 and 1% (v/v) donkey serum. After blocking, cells were incubated for 2 hours at room temperature with primary antibodies diluted in staining solution (PBS supplemented with 0.1% Triton X-100 and 1% donkey serum). The primary antibodies used were: mouse monoclonal anti-T4 (Santa Cruz, sc-52247, ICC 1:50), rabbit polyclonal anti-SLC16A2 (Sigma-Aldrich, HPA003353- 100UL, ICC 1:200), and rabbit polyclonal anti-FLAG (Sigma-Aldrich, F7425-.2MG, ICC 1:100).

Cells were washed with PBS three times for 5 minutes, followed by incubation with Alexa Fluor™ Plus 647 Phalloidin (Thermo Scientific, A30107, ICC 1:400) and Hoechst 33342 (Sigma-Aldrich, B2261, 0.2 μg/ml) diluted in PBS for 30 minutes at room temperature. After incubation, cells were washed with PBS three times for 5 minutes, followed by incubation with secondary antibodies diluted in staining solution for 1 hour at room temperature. The following fluorescence-conjugated secondary antibodies were used: Donkey anti-Rabbit IgG (H+L) Alexa Fluor 488 (Thermo Scientific, A21206, ICC 1:500) and Goat anti-Mouse IgG (H+L) Highly Cross-Adsorbed Alexa Fluor Plus 555 (Thermo Scientific, A32727, ICC 1:500). Cells were washed twice for 5 minutes with PBS. Coverslips were mounted using DAKO Mounting Solution (Agilent DAKO, S302380-2). Confocal microscopy was performed using a Zeiss LSM980 point-scanning confocal microscope based on a Zeiss Observer7 inverted microscope. Images were acquired using a PlanApo 40X/1.4NA oil immersion objective with 488 nm and 561 nm laser lines. All image acquisition was performed using Zen Blue 3.7 software (Zeiss). Image analysis and quantification were carried out with the support of the Human Technopole Image Analysis Facility using python pipelines.

### MCT8 and MCT10 purification

Expi293F™ cells (Thermo Fisher Scientific) were cultured in Dynamis™ AGT™ Medium (Thermo Fisher Scientific) supplemented with 1% (v/v) Pen/Strep (Euroclone) and 4 mM stable glutamine (Euroclone) at 130 rpm, 8% CO2, 37°C, and 85% humidity. These cells were used to produce MCT10, MCT8, and all corresponding mutants. The cells were grown to a concentration of 2-3 x 10^6 cells/mL with a viability greater than 95%. Transfection was performed without a prior media change. For each liter of culture, 1.1 mg of plasmid and 3 mg of polyethyleneimine (PEI Prime, Sigma-Aldrich) were pre-diluted separately in 25 mL of pre-warmed media before being combined. After incubating the plasmid-PEI mixture for 20 minutes at room temperature, the 50 mL complex was added dropwise to the cells. Protein expression proceeded for 72 hours. Cells were then removed by centrifugation at 1,000 rcf for 10 minutes.

The resulting pellet was resuspended in 10 mL of lysis buffer per 1 g of cell mass (25 mM HEPES, pH 7.5, 150 mM NaCl, 1.5% n-dodecyl-β-d-maltopyranoside (βDDM; Anatrace), supplemented with 10 μg/mL DNase I and 1x cOmplete™ EDTA-free protease inhibitor cocktail (Roche). Cell lysis and membrane solubilization was carried out using a Dounce homogenizer and incubated at 4°C for two hours. The soluble fraction was separated via centrifugation (45,000 g for 1 hour). The clarified lysate was then loaded onto 10 mL of M2 anti-Flag affinity resin (Anti-Flag® M2 affinity gel, Sigma-Aldrich), which had been pre-equilibrated with Buffer A (25 mM HEPES, pH 7.5, 150 mM NaCl, 0.04% βDDM). The mixture was gently stirred at 4°C for 1 hour, and the beads were collected by centrifugation at 1,000 g for 10 minutes.

The beads were transferred to a 100 mL EconoColumn (Bio-Rad) and washed with 5 column volumes (CV) of Buffer A. Elution was performed using 5 CV of Buffer A supplemented with 0.2 mg/mL Flag peptide (Tebu bio) to release the MCT8/10 proteins. All fractions were analyzed by SDS-PAGE, and positive fractions were pooled and concentrated to 500 µL using a 100 kDa MW cut-off concentrator (Millipore). Final purification was performed by size-exclusion chromatography on a Superose 6 Increase 10/300 column (GE Healthcare) equilibrated in Buffer A. The fractions containing purified MCT8/10 were combined and concentrated to approximately 1 mg/mL using a 100 kDa MW cut-off concentrator (Millipore), then aliquoted and flash-frozen for storage at -80°C.

### MCT8 and MCT10 wild type Cryo-EM Studies

Single particle cryo-EM was used to ensure the correct folding of MCT8 and MCT10 stabilized in ßDDM. Briefly, 2.5 µL of MCT8 and MCT10 at 4 mg/mL were vitrified in liquid ethane using a Vitrobot Mark IV (FEI). For MCT8, 26166 images were recorded on a Titan Krios G4 transmission electron microscope (Thermo Scientific) operated at 300 kV in EF- TEM mode, and for MCT10, 920 images were recorded on a Glacios transmission electron microscope operated at 200 kV. Both microscopes were equipped with a Falcon 4i (Thermo Scientific) direct electron detector and a Selectris X (Thermo Scientific) post-column energy filter with a slit width of 10 eV. 800000 and 240000 particles were used to generate 2D class averages for MCT8 and MCT10, respectively, using cryoSPARC.

### Anti-ALFATag Nanobody Purification

2xYT medium supplemented with 100 µg/mL ampicillin was inoculated with *E. coli* BL21 (DE3) transformed with pNT1440-HIS-TEVNbALFA. The culture was grown at 37°C with shaking at 200 rpm until OD600 = 0.6. HIS-TEVNbALFA expression was induced by adding 0.55 mM IPTG while decreasing the temperature to 17.5 °C for 16 hours. Cells were harvested by centrifugation at 4500 × g for 20 minutes at 4°C. During purification, the sample was kept at 4°C unless otherwise specified.

Cells were resuspended in cold lysis buffer (50 mM Hepes pH 8, 50 mM NaCl, 5 mM MgCl2 and 1 tablet 50 mL^−1^ EDTA-free protease inhibitor, Roche) plus 0.5 mg/mL lysozyme and 0.1 mg/mL DNase I (Sigma) and stirred at room temperature for 20 minutes. Cells were lysed through three cycles on a CF1 Cell Disruptor at 20kpsi (Constant Systems). The cell lysate was centrifuged at 45,000 × g for 30 minutes, and the insoluble material was collected. The pellet was washed and centrifuged twice with 25 mM Hepes pH 7.5, 150 mM NaCl, 1 M urea, and 1% Triton X-100 before solubilizing the inclusion bodies in 25 mM Hepes pH 7.5, 150 mM NaCl, and 6 M guanidine hydrochloride. HIS-TEVNbALFA was refolded by a overnight dialysis in 25 mM Hepes, 150 mM NaCl, and 10 mM imidazole and clarified by centrifugation at 45,000 × g. The supernatant was incubated for 1 hour with 10 mL of Ni Sepharose High Performance resin (Merck) pre-equilibrated with dialysis buffer. The slurry was washed with dialysis buffer supplemented with 50 mM imidazole before collecting the elution fraction by adding 500 mM imidazole.

The eluted HIS-TEVNbALFA was supplemented with TEVHIS protease in a 1:10 ratio, 1 mM DTT, 0.5 mM EDTA, and incubated for three hours at room temperature with mild shaking, followed by overnight dialysis in 25 mM Hepes and 150 mM NaCl. To remove uncleaved Nanobody and TEVHIS, a reverse IMAC was performed and the flowthrough collected. The sample was concentrated using an Amicon® Ultra (MWCO 3 kDa) up to 5 mL and injected on a Superdex75 16/600 PrepGrade column (Cytiva). The eluted sample was stored at -80°C for future use.

### Sample Preparation for Cryo-EM Structural Studies

Purified MCT8/10 constructs bearing the alfa-tag were incubated with a 3-fold molar excess of nanobody α-alfa tag for 1 hour at 4°C with gentle shaking. A ten-fold excess (mg: mg) of Amphipols (PMAL-C8, Anatrace) was then added and incubated overnight. The detergent was removed by incubating the sample with Biobeads (Bio-Rad) for 2 hours at 4°C. Final polishing was achieved by injecting the sample onto a Superdex 200 10/300 column equilibrated with 25 mM Hepes, pH 7.5, and 150 mM NaCl. For vitrification, 2.5 µL of a 0.5 mg/mL sample, optionally supplemented with 20 µM T4 or 40 µM Silychristin, was incubated for 10 seconds onto an R1.2/1.3 Quantifoil 200 mesh copper grid (Electron Microscopy Science) before vitrification in liquid ethane using a Vitrobot Mark IV (FEI).

### Cryo-EM Data Collection

Data acquisition was performed on a Titan Krios G4 transmission electron microscope (Thermo Scientific) operated at 300 kV in EF-TEM mode and equipped with a Falcon 4i (Thermo Scientific) direct electron detector and a Selectris X (Thermo Scientific) post-column energy filter with a slit width of 10 eV. All datasets were collected using EPU automated acquisition software in electron counting mode at a nominal magnification of 165,000x, corresponding to a calibrated pixel size of 0.748 Å/pixel, and at defocus range of -0.8 to -2.0 µm. The datasets were recorded in Electron Event Representation (EER) file format. A total of 44,327 movies were collected for MCT8-D424N, 36,940 for MCT8- Silychristin, 41,224 for MCT8, 22,504 for MCT8-T4, and 36,610 for MCT10-T4. EER files were fractionated to achieve a dose of 1 e- Å-2 per frame.

### Cryo-EM Data Processing

The processing workflow is illustrated in Supplementary Fig. 2. EER movies were averaged using MotionCorr within Relion5^25,26^, and the corrected micrographs were exported to CryoSPARC^27^. Contrast Transfer Function (CTF) parameters were calculated using Patch CTF estimation. For all datasets, approximately 7-10 million particles were extracted and sorted through at least five iterations of 2D class average classifications, yielding about 500,000 to 800,000 particles. This particle stack was further classified through a series of Ab-initio reconstructions (two or three classes) followed by Non-uniform refinement and local refinement iterative cycles until resolution, B-factor, and Fourier space completeness ceased to improve. The resulting stack of approximately 100,000 - 180,000 particles was imported into Relion5 for 3D Classification (optional) and Auto-refine with blush enabled. CTF refinement and Bayesian polishing were performed on the final particle stack, and amphipol densities were removed during post-processing using a mask generated from the PDB model.

### Model building

For MCT8, an AlphaFold2 inward-facing model template was downloaded from UniProt (P36021). The outward-facing MCT10 template model was generated using Swiss-Model^28^ with MCT1 (PDB ID: 7YR5)^22^ as the input. For both MCT8 and MCT10, the models were rigid-body fitted into the cryo-EM maps and refined using Coot^29^. Further refinement was performed with iSOLDE^30^ and Phenix^31^. Ligand restraints for T4 and silychristin were generated using Elbow. Further information available in Table 1. Model analysis and visualization was performed in PyMol and ChimeraX ^32,33^. Conservation analysis was performed with ConSurf^34^.In most scientific literature on MCT8 mutations the position of the mutations is indicated using the NM_006517.3 reference sequence and uses +1 as the A of the ATG translation initiation codon of the long MCT8 isoform, with the initiation codon as codon 1. To facilitate interpretation, we added the position of residues in the long isoform between brackets in the legends of relevant figures.

### Microscale Thermophoresis

MicroScale Thermophoresis (MST) MCT8/10 (wt and mutants) were tested for interaction with ligands of interest using a Monolith instrument (Nanotemper) using LabelFree standard capillaries, monitoring intrinsic protein fluorescence. All the ligands (T4,T3, TRIAC, Silychristin) were resuspended in DMSO, and a 16-point dilution series was prepared over specific concentration range in the final assay buffer (25 mM Hepes pH 7.5, 150 mM NaCl, 0.04 % bDDM and 5 % DMSO). Binding reactions were prepared adding 50 nM of protein incubating 10 minutes at room temperature. Since addition of ligands induced decrease in fluorescence signal, an SDS-denaturation (SD) test was performed, following the manufacturer protocol, to ensure fluorescence changes were not caused by non-specific effects. Since in denaturing conditions the fluorescent signal of the ligand-containing points was unchanged with respect to that of ligand-free points, we concluded that initial fluorescence intensities might be used to ascertain ligand binding and for equilibrium dissociation constants (K*D*) calculation. Measurements were conducted with 100 % excitation power and MST power set to high, initial fluorescence data were analyzed, and K*D* were obtained using GraphPad Prism. For the competitive binding experiments, MCT8/10 was incubated overnight with either 100 µM T4 or 50 µM Silychrstin. All experiments were performed in biological triplicates.

### Isothermal Titration Calorimetry (ITC)

The thermodynamics of the MCT8-Silychristin interaction were determined using Isothermal Titration Calorimetry (ITC) on a MicroCal PEAQ ITC instrument (Malvern Panalytical). A 10 µM MCT8 solution was titrated with Silychristin (100 µM) using injection of 2 µL. Protein and ligand were prepared and diluted to be in the same final buffer (25 mM Hepes pH 7.5, 150 mM NaCl, 0.04 % bDDM). Calorimetric data were analyzed with the instrument software, providing thermodynamic fitting parameters such as the reaction enthalpy change (ΔH, cal mol⁻¹) and the binding constant (Ka, M⁻¹). The reaction entropy (ΔS) was derived using the equations ΔG = −RTlnKa (where R = 1.9872 cal mol⁻¹ K⁻¹, and T = 298 K) and ΔG = ΔH − TΔS. The best fit was determined using the statistical goodness of fit (GoF) parameter. The measurement was repeated three times, and after each run, the sample was collected and injected onto a Superose 6 10/300 column to assess the dispersity.

## Acknowledgements

We thank all members of Human Technopole facilities and in particular: Paolo Swuec, Gaetano D’Urso, Simona Sorrentino of the Cryo-Electron Microscopy Unit of the National Facility for Structural Biology for technical support and assistance with Cryo-EM sample preparation and data collection, the bioimage analysis infrastructure unit of the national facility for data handling and analysis, the National Facility for Light Imaging for the assistance with image acquisition, the Biomass production Unit, the National Facility for Structural Biology – IU3 Biophysics Unit. We thank Daniele Colombo from the IT department for pivotal help with software installation and debugging, Eliana Bianco for methodological support on cell-based assays and Karthik Kévin Ramanadane for useful comments on the work. We acknowledge funding from Human Technopole and from the European Research Council (ERC-2021-STG Thyromol #101041298) to FC and by Eurostars (project number E11337) to WEV.).

## Data availability

The data that support this study are available from the correspondence upon request. Atomic coordinates have been deposited in the Protein Data Bank (PDB) and Electron microscopy Data Bank under accession codes 9GSZ / EMD-51559 (Human monocarboxylate 10 bound to L-thyroxine), 9GF8 / EMD-51311 (Human Monocarboxylate Transporter 8), 9FOT / EMD-50629 (Human monocarboxylate transporter 8 bound to Silychristin), 9FKN / EMD-50523 (Human monocarboxylate transporter 8 bound to L-thyroxine), 9GV5, EMD-51624 (Human monocarboxylate transporter 8 D424N mutant). Source data is provided in this paper.

## Author contributions

MT performed clonings, biochemistry, biophysics, EM data collection and processing, model building and structural analysis

GT performed transport and inhibition experiments in cells

IB performed preliminary biochemistry and structural analysis of MCT8

FM performed clonings, biochemistry and biophysics experiments assisted by MT SP provided help with biophysics data collection and analysis

MF and CS performed cloning.

SG, FSvG, WEV, FC, MT, GT provided interpretation to the data

FC, WEV wrote the draft manuscript. All authors assisted with manuscript preparation.

