## Supplementary Information for "Molecular mechanism of thyroxine transport by monocarboxylate transporters"

#### Supplementary Figure 1

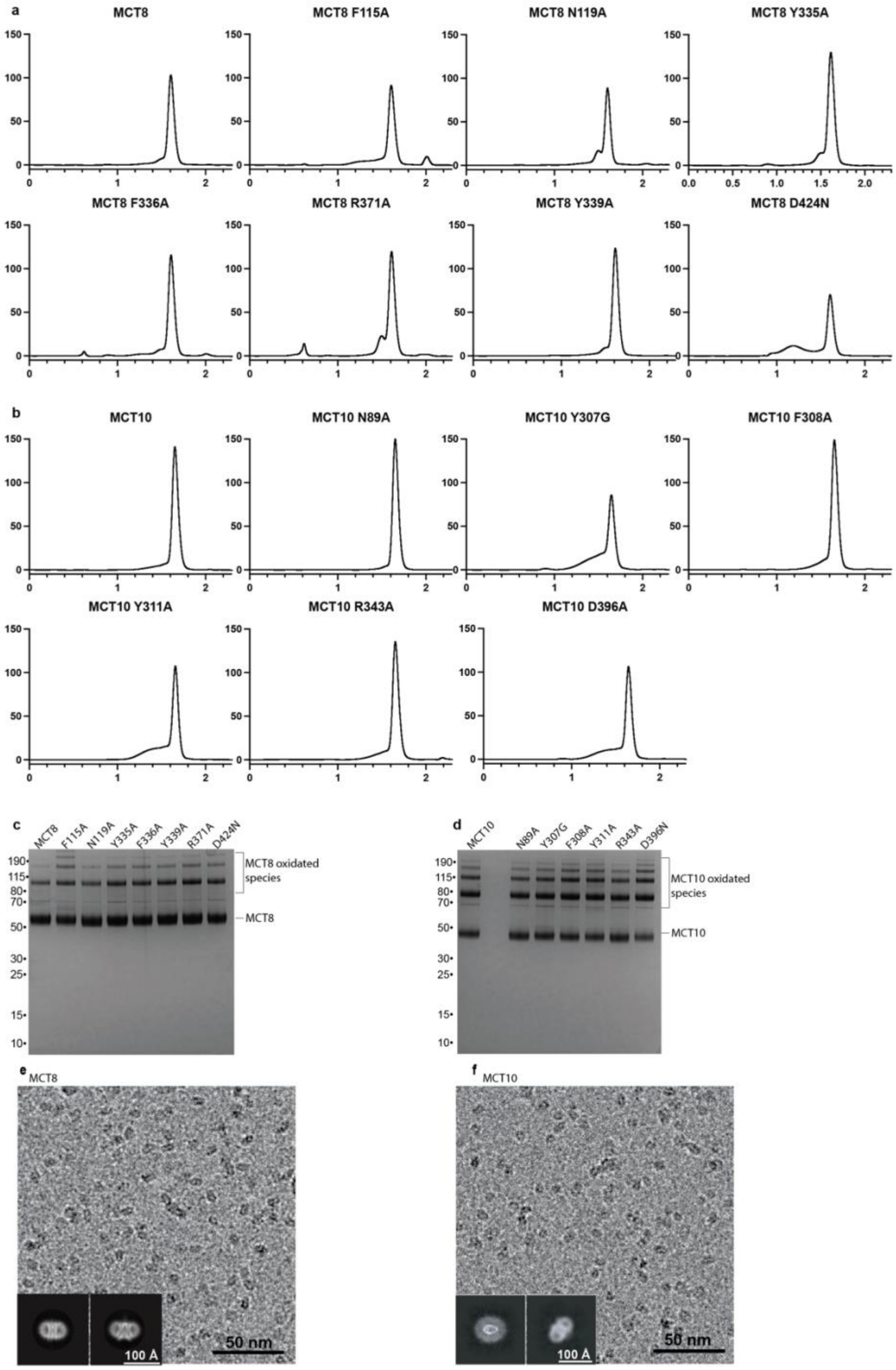


**Supplementary Figure 1:** Protein purification of MCT variants in detergent. Size exclusion chromatography profiles of MCT8 (a) and MCT10 (b) performed in bDDM detergent. c,d. SDS-PAGE of MCT8 and MCT10 variants. e-f. Representative cryo-EM micrographs and 2D class averages showing folding of WT MCT8 and MCT10 in ßDDM.

#### Supplementary Figure 2

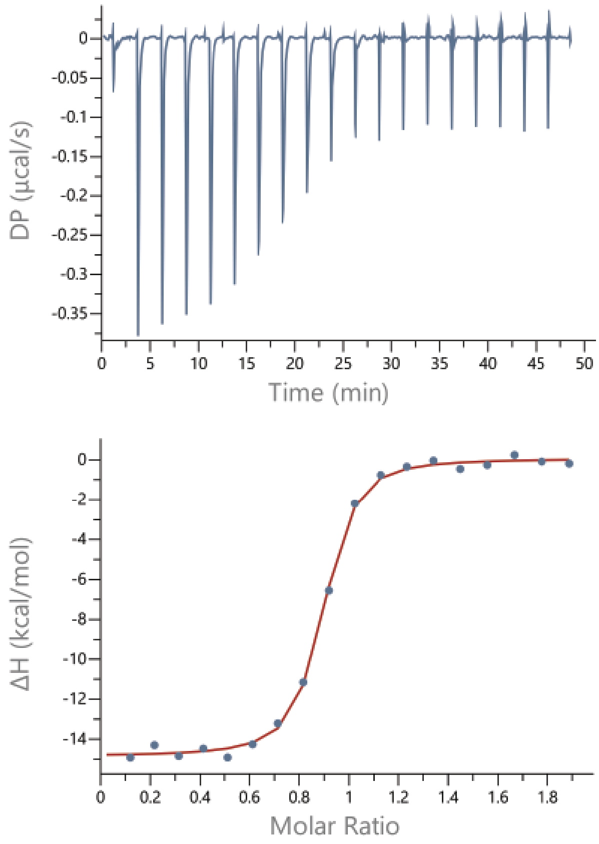


**Supplementary Figure 2:** Binding of Silychristin to MCT8 measured by Isothermal titration calorimetry (ITC) in ßDDM. Kd : 44.5 nM ±6

#### Supplementary Figure 3

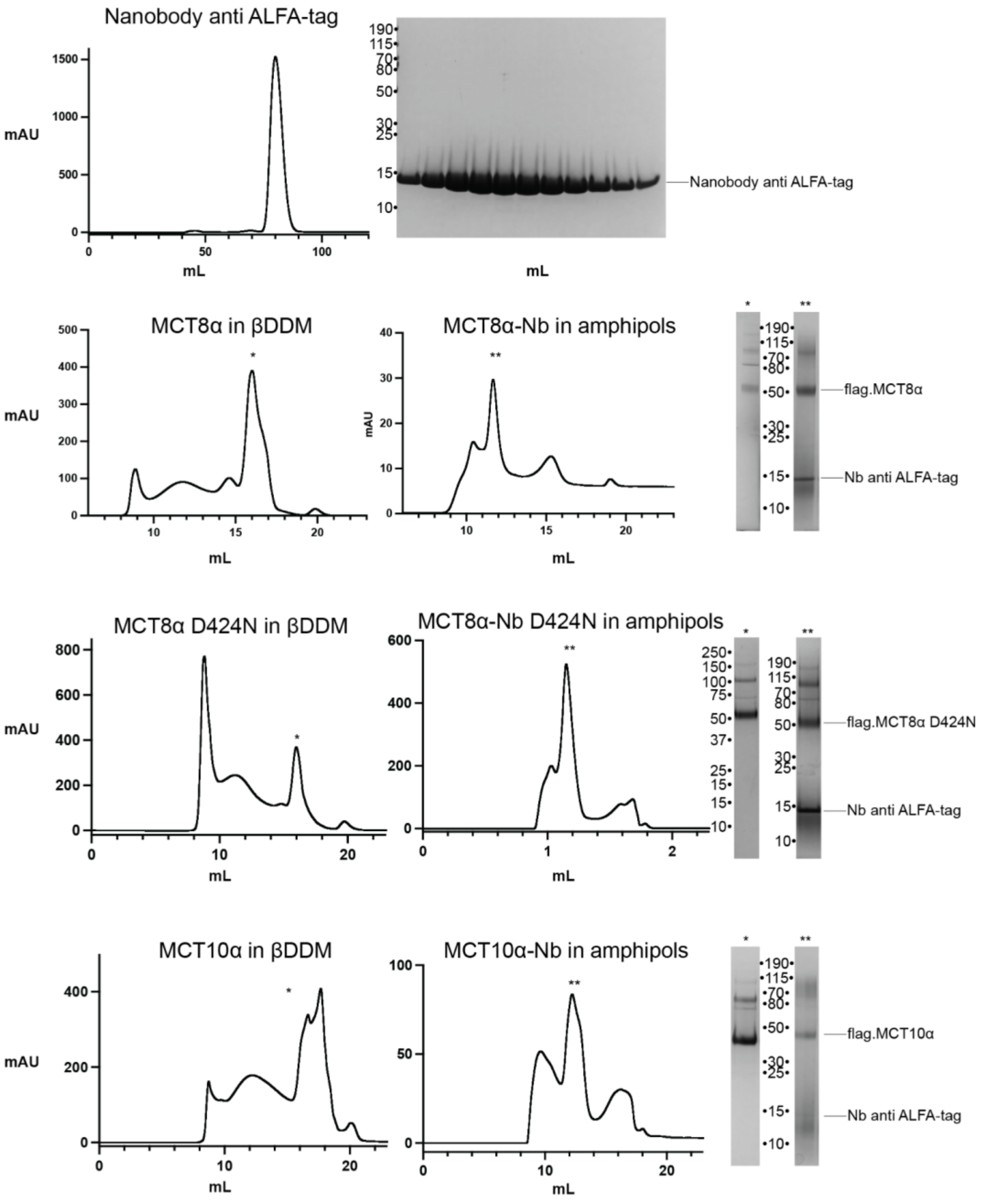


**Supplementary Figure 3:** Purification of MCT-ALFA/NbALFA complexes in presence of amphipols used for cryo-EM structural analysis (Size exclusion chromatography and relative SDS-PAGE are reported)

#### Supplementary Figure 4

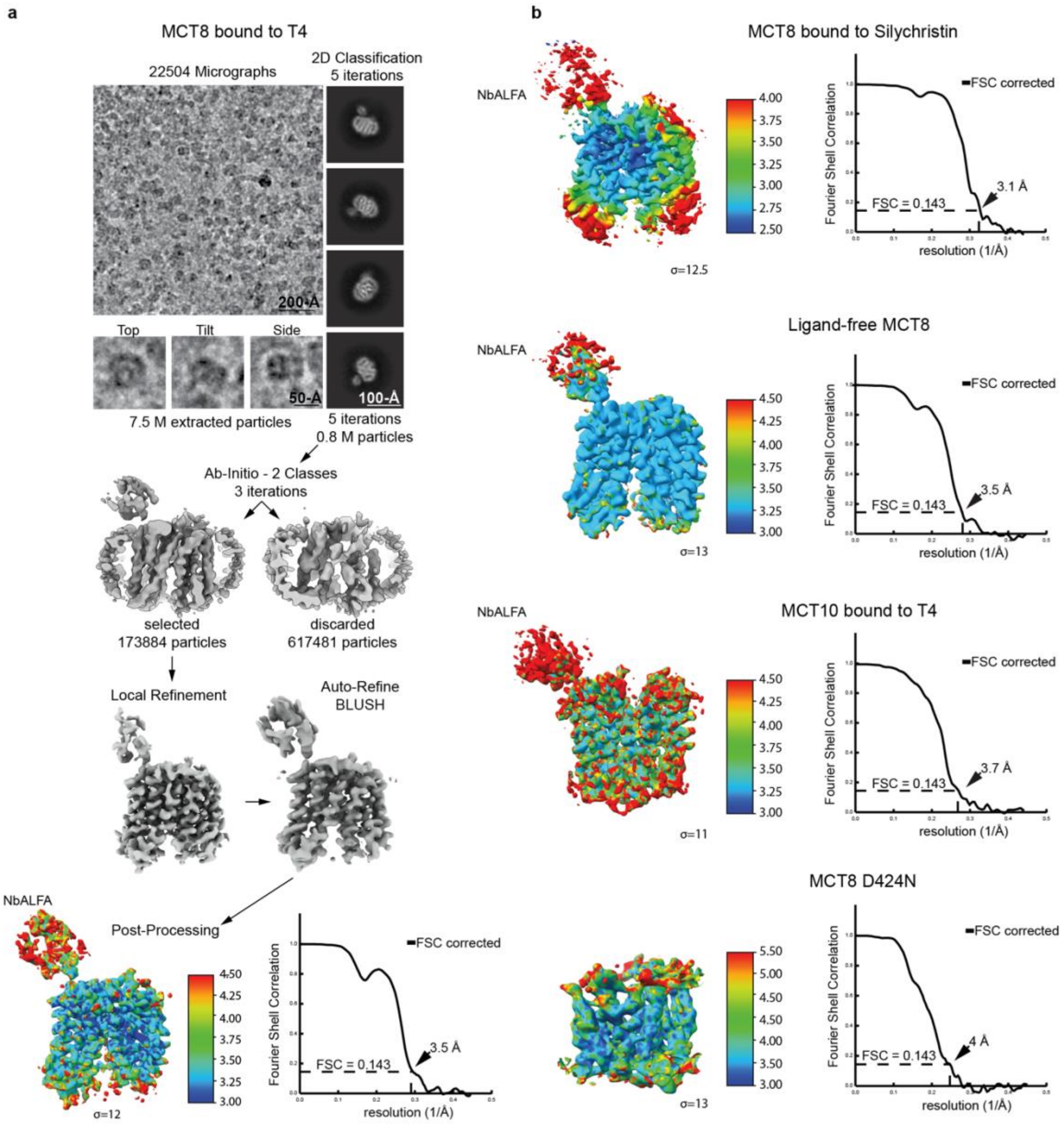


**Supplementary Figure 4: a.** Cryo-EM image processing workflow example for MCT8-T4 map **b.** Resulting maps for MCT8-Silychristin, ligand-free MCT8, D424N mutant coloured by local resolution with relative FSC curve.

#### Supplementary Figure 5

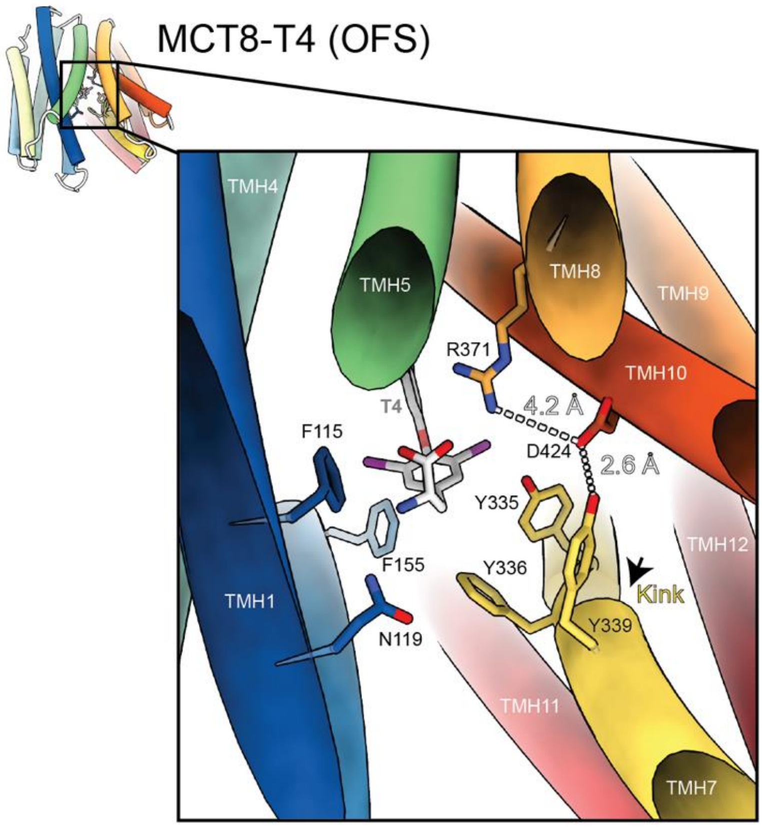


**Supplementary Figure 5:** Closed-up view of T4-bound MCT8 structure in the outward-facing state (OFS) showing interactions between D424 (TMH10) Y339 (TMH7) and R371 (TMH8) and spatial relationship to the substrate.

#### Supplementary Figure 6

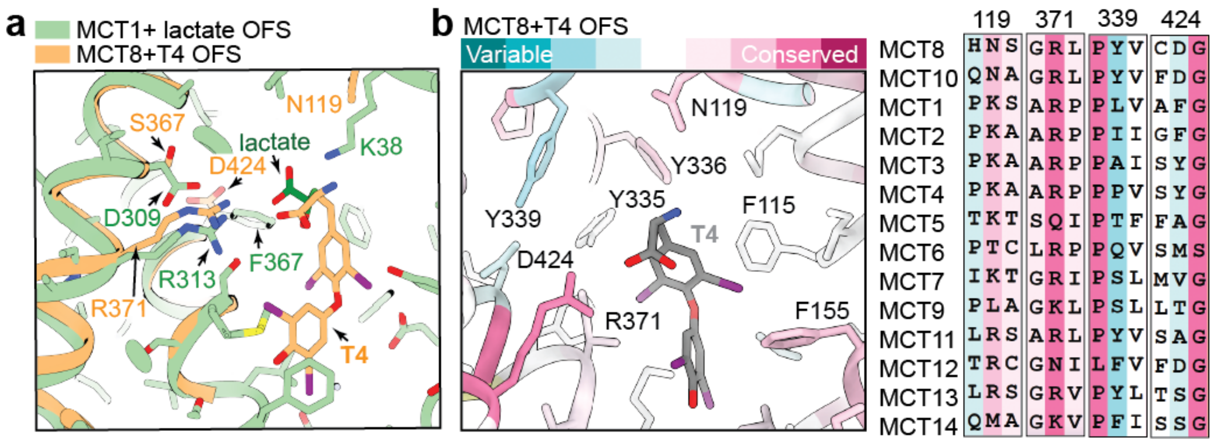


**Supplementary Figure 6: Structural comparison between monocarboxylate transporters. a.** Overlap between MCT8-T4 OFS and the structure of MCT1 PDB ID: 6LZ0 in complex with lactate (OFS). The carboxylic group of T4 is very close to the lactate position and it is coordinated by the conserved R371/313, whereas other residues are variable **b.** Sequence conservation (Consurf) of residues in the substrate binding pocket for all MCTs. While R371 is highly conserved, the residues characteristic MCT8 and MCT10 are the pair Y339 and D424. To derive the position of residues in the long isoform of MCT8, 74 amino acids should be added (between brackets in the legend).

#### Supplementary Figure 7

| Construct | Primers |
|---|---|
| Flag-tag Insertion in pcDNA3_hMCT10 | FW_AAAAAGCTTATGGATTACAAGGATGACGACGATAAGATGGTGCTCTCCCAGGAGGAG |
| | RV_AAATCTAGATTAAATAATAGAGTCAGATTC |
| pcDNA3_hMCT10_N89A | FW_ATCCAGGCCGCTTGCGGGGTG |
| | RV_CAAGCGGCCTGGATGCCGAACAC |
| pcDNA3_hMCT10_Y307G | FW_TTGGAGGCTTTGTGCCTTATGTTCACTTG |
| | RV_AGGCACAAAGGCTCCAAAAAGTGCAAG |
| pcDNA3_hMCT10_F308A | FW_GGATACGCTGTGCCTTATGTTCACTTGATG |
| | RV_AGGCACAGCGTATCCAAAAAGTGCAA |
| pcDNA3_hMCT10_Y311A | FW_GCCTGCTGTTCACTTGATGAAACATG |
| | RV_GTGAACAGCAGGCACAAAGTATCCA |
| pcDNA3_hMCT10_R343A | FW_GTTGGAGCACTGCTCTTTGGCC |
| | RV_AGCAGTGCTCCAACTCCTGAAGTGA |
| pcDNA3_hMCT10_D396A | FW_CTCTTCGCTGGATGCTTCATTTCCATTATGG |
| | RV_GCATCCAGCGAAGAGACCCATGATG |
| pcDNA3_hMCT10_alfa-tag_P470 | FW_CGAGGAGGAACTAAGAAGGCGACTAACAGAACCATAATCTAGAGGGCCCGTTTAAAC CCGC |
| | RV_CTTCTTAGTTCCTCCTCGAGTCTGCTCGGGATAAAACAAAGCACAGCACCTCC |
| pcDNA3_hMCT8_F115A | FW_GCTCCATCGCCGGCATCCATAACTCTGTCGGGATCCTCTACTCCATG |
| | RV_ATGCCGGCGATGGAGCCGTTGCACCAGGTGGCAG |
| pcDNA3_hMCT8_N119A | FW_GGCATCCATGCCTCTGTCGGGATCCTCTACTCCATGC |
| | RV_GACAGAGGCATGGATGCCGAAGATGGAGCCG |
| pcDNA3_hMCT8_Y335A | FW_CCCTTGGCGCCTTTGTTCCCTATGTACACCTGATGAAGTATGTGGAGG |
| | RV_ACAAAGGCGCCAAGGGCAGCAGCAGC |
| pcDNA3_hMCT8_F336A | FW_CTTGGCTACGCAGTTCCCTATGTACACCTGATGAAGTATGTGGAGGA |
| | RV_GGGAACTGCGTAGCCAAGGGCAGCAGCAGCAATTCCGAAGGCCCAG |
| pcDNA3_hMCT8_Y339A | FW_CTTTGTTCCCGCAGTACACCTGATGAAGTATGTGGAGGAGGAGT |
| | RV_GGTGTACTGCGGGAACAAAGTAGCCAAGGGCAGC |
| pcDNA3_hMCT8_R371A | FW_GCCTTGGGGCACTTGTGTCAGGCCACATCAGTGAC |
| | RV_CACAAGTGCCCCAAGGCCTGAGGTAGCCC |
| pcDNA3_hMCT8_D424A | FW_CCTTTGCGCAGGCTTCTTCATCACCATCATGGCCC |
| | RV_AGCCTGCGCAAAGGCCCAGGAAAAGACAGACG |
| pcDNA3_hMCT8_alfa-tag_P505 | FW_CGAGGAGGAACTAAGAAGGCGACTAACAGAACCAATCTAATCTAGAGGGCC |
| | RV_GCCTTCTTAGTTCCTCCTCGAGTCTGCTAGGGACGAAGAAGAGGATTACAGC |
| pcDNA3_hMCT8_alfa-tag_F512 | FW_CGAGGAGGAACTAAGAAGGCGACTAACAGAACCAATCTAATCTAGAGGGCC |
| | RV_GCCTTCTTAGTTCCTCCTCGAGTCTGCTAGGGAACATCCTTTGATGCATCAGAGG |

**Supplementary Figure 7:** DNA primers used for the cloning of MCT8 and MCT10 constructs

#### Supplementary Figure 8

flag.MCT8 alfa-tag P505
MDYKDDDDKALQSQASEEAKGPWQEADQEQQEPVGSPEPESEPEPEPEPEPVPV
PPPEPQPEPQPLPDPAPLPELEFESERVHEPEPTPTVETRGTARGFQPPEGGFGW
VVVFAATWCNGSIFGIHNSVGILYSMLLEEEKEKNRQVEFQAAWVGALAMGMIFFC
SPIVSIFTDRLGCRITATAGAAVAFIGLHTSSFTSSLSLRYFTYGILFGCGCSFAFQPS
LVILGHYFQRRLGLANGVVSAGSSIFSMSFPFLIRMLGDKIKLAQTFQVLSTFMFVLM
LLSLTYRPLLPSSQDTPSKRGVRTLHQRFLAQLRKYFNMRVFRQRTYRIWAFGIAA
AALGYFVPYVHLMKYVEEEFSEIKETWVLLVCIGATSGLGRLVSGHISDSIPGLKKIY
LQVLSFLLLGLMSMMIPLCRDFGGLIVVCLFLGLCDGFFITIMAPIAFELVGPMQASQ
AIGYLLGMMALPMIAGPPIAGLLRNCFGDYHVAFYFAGVPPIIGAVILFFVPSRLEEEL
RRRLTEPI*

flag.MCT8 D424N alfa-tag F512
MDYKDDDDKALQSQASEEAKGPWQEADQEQQEPVGSPEPESEPEPEPEPEPVPV
PPPEPQPEPQPLPDPAPLPELEFESERVHEPEPTPTVETRGTARGFQPPEGGFGW
VVVFAATWCNGSIFGIHNSVGILYSMLLEEEKEKNRQVEFQAAWVGALAMGMIFFC
SPIVSIFTDRLGCRITATAGAAVAFIGLHTSSFTSSLSLRYFTYGILFGCGCSFAFQPS
LVILGHYFQRRLGLANGVVSAGSSIFSMSFPFLIRMLGDKIKLAQTFQVLSTFMFVLM
LLSLTYRPLLPSSQDTPSKRGVRTLHQRFLAQLRKYFNMRVFRQRTYRIWAFGIAA
AALGYFVPYVHLMKYVEEEFSEIKETWVLLVCIGATSGLGRLVSGHISDSIPGLKKIY
LQVLSFLLLGLMSMMIPLCRDFGGLIVVCLFLGLCNGFFITIMAPIAFELVGPMQASQ
AIGYLLGMMALPMIAGPPIAGLLRNCFGDYHVAFYFAGVPPIIGAVILFFVPLMHQRM
FPSRLEEELRRRLTEPI*

flag.MCT10 alfa-tag P470
MDYKDDDDKMVLSQEEPDSARGTSEAQPLGPAPTGAAPPPGPGPSDSPEAAVEK
VEVELAGPATAEPHEPPEPPEGGWGWLVMLAAMWCNGSVFGIQNACGVLFVSML
ETFGSKDDDKMVFKTAWVGSLSMGMIFFCCPIVSVFTDLFGCRKTAVVGAAVGFV
GLMSSSFVSSIEPLYLTYGIIFACGCSFAYQPSLVILGHYFKKRLGLVNGIVTAGSSVF
TILLPLLLRVLIDSVGLFYTLRVLCIFMFVLFLAGFTYRPLATSTKDKESGGSGSSLFS
RKKFSPPKKIFNFAIFKVTAYAVWAVGIPLALFGYFVPYVHLMKHVNERFQDEKNKE
VVLMCIGVTSGVGRLLFGRIADYVPGVKKVYLQVLSFFFIGLMSMMIPLCSIFGALIA
VCLIMGLFDGCFISIMAPIAFELVGAQDVSQAIGFLLGFMSIPMTVGPPIAGLLRDKLG
SYDVAFYLAGVPPLIGGAVLCFIPSRLEEELRRRLTEP*

**Supplementary Figure 8:** Protein sequences of purified constructs, alfa tag in purple, flag-tag in green and point mutation in red

### Supplementary Table 1

| Data collection | MCT8 | MCT8-T4 | MCT8-silychristin | MCT10-T4 | MCT8 D424N |
|---|---|---|---|---|---|
| Electron microscope | Titan Krios | Titan Krios | Titan Krios | Titan Krios | Titan Krios |
| Voltage (kV) | 300 | 300 | 300 | 300 | 300 |
| Pixel size (Å) | 0.748 | 0.748 | 0.748 | 0.748 | 0.748 |
| Electron exposure (e−/Å$^2$) | 70 | 70 | 70 | 70 | 70 |
| Defocus range (μm) | 0.8-2.0 | 0.8-2.0 | 0.8-2.0 | 0.8-2.0 | 0.8-2.0 |
| Images | 41224 | 22504 | 36940 | 36610 | 44327 |
| **3D reconstruction** | | | | | |
| Final particle | 109595 | 173884 | 101856 | 57707 | 52344 |
| Resolution (Å) | 3.536 | 3.483 | 3.08 | 3.704 | 4 |
| FSC threshold | 0.143 | 0.143 | 0.143 | 0.143 | 0.143 |
| B factor (Å$^2$) | -124.383 | -75 | -89 | -131.613 | -70 |
| **Refinement** | | | | | |
| Model | MCT8-NbALFA | MCT8-NbALFA bound to thyroxine | MCT8-NbALFA bound to silychristin | MCT10 bound to thyroxine | MCT8 D424N |
| Chains | 2 | 2 | 2 | 1 | 1 |
| **Model Composition** | | | | | |
| Total atoms | 7897 (Hydrogens: 3969) | 7939 (Hydrogens: 3984) | 8008 (Hydrogens: 4014) | 6032 (Hydrogens: 3064) | 5465 (Hydrogens: 2766) |
| Total residues | 508 | 508 | 511 | 383 | 355 |
| Ligand | | T44 | SLT | T44 | |
| **R.m.s deviations** | | | | | |
| Bond length (Å) (# > 4σ) | 0.002 (0) | 0.003 (0) | 0.004 (0) | 0.003 (0) | 0.002 (0) |
| Angles (°) (# > 4σ) | 0.422 (2) | 0.473 (2) | 0.522 (3) | 0.547 (1) | 0.520 (1 |
| **Validation** | | | | | |
| MolProbity score | 1.35 | 1.26 | 1.29 | 1.78 | 1.71 |
| Clashscore | 4.18 | 2.02 | 3.01 | 4.98 | 5.13 |
| Poor rotamers (%) | 0.96 | 0.24 | 0.24 | 1.26 | 0.35 |
| **Ramachandran plot (%)** | | | | | |
| Outliers | 0.00 | 0.00 | 0.00 | 0.00 | 0.00 |
| Allowed | 2.80 | 4.20 | 3.18 | 6.90 | 6.88 |
| Favoured | 97.20 | 95.80 | 96.82 | 93.10 | 93.12 |
| **B-factors/ADP protein** | | | | | |
| Minimum | 48.85 | 5.12 | 26.62 | 30.00 | 89.08 |
| Maximum | 144.99 | 89.95 | 174.66 | 105.95 | 182.99 |
| Mean | 91.34 | 41.73 | 63.63 | 67.47 | 125.92 |
| **B-factors/ADP ligand** | | | | | |
| Minimum | | 23.98 | 42.60 | 23.98 | |
| Maximum | | 63.16 | 42.60 | 63.16 | |
| Mean | | 45.14 | 42.60 | 45.14 | |
| **Model vs. Data** | | | | | |
| CC (mask) | 0.65 | 0.65 | 0.81 | 0.73 | 0.61 |
| CC (volume) | 0.64 | 0.67 | 0.78 | 0.71 | 0.62 |
| Mean CC for ligands | | 0.66 | 0.79 | 0.50 | |

**Supplementary Table 1:** Cryo-EM data collection, refinement and validation statistics
